# Combining transcriptomic and fitness data reveals additive and mostly adaptive plastic responses of gene expression to multiple stress in *Tribolium castaneum*

**DOI:** 10.1101/442145

**Authors:** Eva L. Koch, Frédéric Guillaume

## Abstract

Gene expression is known to be highly responsive to the environment and to vary between species or populations under divergent selection. Yet, its contribution to the process of adaption is still controversial despite growing evidence that differences in gene regulation contribute to adaptive divergence. While most studies so far investigated evolved plasticity in already diverged populations, phenotypic selection acting on gene expression at the onset of adaptation to an environmental change has not been characterized. Here, we combined fitness and whole-transcriptome data in a large-scale experiment with *Tribolium castaneum* to investigate gene expression and fitness responses to drought, heat and their combination. Fitness was reduced by both stressors and their combined effects were nearly additive. Accordingly, expression data showed that both stressors were acting independently and did not interfere physiologically. With expression and fitness within the same individuals, we estimated selection on single gene expression levels. We found that variation in fitness can be attributed to gene expression variation. Selection intensities on expression levels differed between conditions and were opposite between control and stress conditions, showing evidence of evolutionary trade-offs. Plastic expression changes were mostly adaptive when affected by heat stress, and partially non-adaptive when affected by drought.

## Introduction

One of the major goals of evolutionary biology is to understand the genetic basis of phenotypic variation and how it is shaped by natural selection. The mapping of genetic to phenotypic variation depends on many cellular processes, of which RNA transcript abundance, or gene expression, has been shown to play a central role (Abzhanov et al. 2004; Prud’ homme et al. 2007; Carroll 2008; Fay and Wittkopp 2008; Romero et al. 2012). For variation in expression to be relevant for evolution it needs a heritable genetic basis and a link with fitness variation. While the heritability of expression variation has been established in many cases (Cui et al. 2006; Ayroles et al. 2009; Skelly et al. 2009; Long et al. 2012; Morris et al. 2014; McCairns et al. 2016), we still lack direct estimates of the strength of selection acting on transcript level abundance. The link between expression levels and fitness variation is not obvious, since mRNA abundance must be translated into protein abundance, enzyme activity and ultimately phenotypic variation (Feder and Walser 2005; Evans 2015). So far, the evidence for a link between fitness and gene expression variation is mixed and mostly indirect. For instance, in yeast, the knocking-out of many genes had inconsequential effects on fitness (Giaever et al. 2002), whereas more recent evidence showed that variation in expression can significantly affect fitness (Keren et al. 2016; Sato et al. 2016). Unfortunately, data in more complex organisms are still scarce, especially on a transcriptome-wide scale. Indirect evidence supporting the importance of gene expression in evolution comes from studies showing differences in expression levels between adaptively divergent populations in yeast (Townsend et al. 2003), humans (Fraser 2013), *Drosophila* (Hutter et al. 2008) or fish (Whitehead and Crawford 2006; McCairns and Bernatchez 2009; Morris et al. 2014; Dayan et al. 2015; Ghalambor et al. 2015; Leder et al. 2015). In such cases, further support can be brought when evolved differences in regulatory DNA sequences are found (Zheng et al. 2011; Sato et al. 2016). Additionally, experimental evolution approaches were successful to detect altered expression levels within a few generations in multiple organisms that adapted to different environmental conditions (Riehle et al. 2003; Telonis-Scott et al. 2009; Yampolsky et al. 2012; Ghalambor et al. 2015; Huang and Agrawal 2016).

Beside genetic divergence causing expression differentiation (Brem et al. 2002; Fay and Wittkopp 2008; Romero et al. 2012), phenotypic plasticity can also play an important role in differentiation, especially at the onset of adaptation to novel environments (Yeh and Price 2004; Lande 2009; Chevin et al. 2010; Ehrenreich and Pfennig 2016). Expression is a highly plastic trait and is involved in the immediate response of organisms to changes in their environment (Gasch et al. 2000; Chen et al. 2003; Gibson 2008; López-Maury et al. 2008; Des Marais et al. 2013). The role of plasticity in evolution is, however, contentious. It is often argued that if plasticity is adaptive, it should impede evolution since it can hide genetic variance on which selection would act, weakening selection (Price et al. 2003). Yet, plasticity is also key for population persistence in a changing environment because it can keep populations at higher sizes, or buffer novel variants against purifying selection (Chevin et al. 2010; Fitzpatrick 2012), thus facilitating long-term adaptation (Yeh and Price 2004; Lande 2009; Ehrenreich and Pfennig 2016). Studies comparing plastic and evolved responses of gene expression in natural populations repeatedly found that plasticity was non-adaptive because opposed in direction to the evolutionary response (Sørensen et al. 2001; Dayan et al. 2015; Ghalambor et al. 2015). It may thus be that non-adaptive plasticity facilitates evolutionary divergence by increasing the strength of selection (Ghalambor et al. 2007, 2015).

Unfortunately, we currently lack estimates of fitness effects of expression changes during early stages of adaptation, on which depends the adaptive value of those changes. Current studies have inferred the adaptive value of a plastic response of expression by comparing the direction of change of differentially expressed genes between evolved and non-evolved populations or experimental lines in common gardens. An early-stage expression change in the same direction as an evolved change (e.g., when comparing ancestral to evolved lines) would be interpreted as adaptive, without knowledge of the immediate fitness effects of the early-stage expression change (Whitehead and Crawford 2006; McCairns and Bernatchez 2009; Morris et al. 2014; Ghalambor et al. 2015; Leder et al. 2015). However, the two may differ because plasticity can evolve during adaptation (e.g., due to canalization, Heckel et al. 2016), or because beneficial immediate stress responses can be costly to maintain (Moya et al. 2012, 2015), and thus reverted during long-term adaptation (Fong et al. 2005; Rodŕiguez-Verdugo et al. 2016; Ho and Zhang 2018). Evolutionary changes in plasticity make post-adaptation inferences of the adaptive value of early-stage expression changes uncertain. In contrast, measuring fitness directly in organisms that have been exposed to a recent change in conditions can give us deeper insight into the initial processes leading to adaptive divergence between populations. Knowing the strength of selection acting on early-stage plastic transcriptional responses can tell us more about the immediate adaptive value of plasticity in gene expression and the targets of selection during the early stage of adaptation. Ultimately, it can also tell us how short-term selection pressures are linked to long-term optimum expression levels.

The difference between short and long-term gene expression changes will depend on trade-offs between the benefits of immediate stress responses and their long-term costs. Adaptation necessitates optimal re-allocation of energy resources between maintenance and reproduction. However, optimal solutions for this trade-off may differ between environmental stressors (Schlichting and Smith 2002), which further complicates the study of plastic responses in gene expression and their fitness effects in variable environments. A beneficial response elicited by one environmental factor may be overridden by a negative effect in presence of a second factor and generate a pattern of non-adaptive plasticity. The joint effects of stress factors may not be simply deduced from single responses and may result in complex interactions when in combination (Crain et al. 2008; Holmstrup et al. 2010; Byrne and Przeslawski 2013; Gobler et al. 2014; Gunderson et al. 2016). It is thus crucial to understand the trade-offs faced by organisms when adapting to changed environments (DeBiasse and Kelly 2016; Kelly et al. 2016). Transcriptomics can give us insights into the mechanisms underlying trade-offs between 1) responses to different stressors, and 2) energy allocation trade-off between reproduction and maintenance within conditions. It can potentially show which physiological processes are activated, thereby giving us information about how resources are used. Trade-offs in stress responses can not only limit plastic responses but may also constrain evolution and future adaptation (Etterson and Shaw 2001; Pörtner et al. 2006; Kelly et al. 2016). Therefore, linking the immediate plastic responses of gene expression with their fitness effects at the onset of adaptive divergence should allow a better understanding of their adaptive value and evolutionary consequences.

In this study we asked how *Tribolium castaneum* (the red flour beetle) was affected by heat and drought in single stressor treatments and in a combination treatment. We combined a fitness assay with measuring gene expression using RNA-seq. This allowed us to get more insight into the physiological processes affected by different stressors, to examine whether the transcriptomic responses to heat and drought overlapped and if gene expression was changed in the same direction by both stressors. Based on this, we can make predictions about how stressors will act in combination and test this in the combined stressor treatment. We further examined the potential resource allocation trade-off between reproduction and stress response by focusing on specific genes known to be involved in reproduction. We tested how these genes were affected by the different treatments, and whether this corresponded with the observed fitness consequences. Since we measured expression and fitness in the same individuals in sufficient sample size, we were also able to estimate transcriptome-wide selection intensities on gene expression levels giving us an unprecedented view of selection pressures on gene expression in different environments. Further, we were able to examine whether immediate plastic responses were adaptive or non-adaptive, thus contributing to our understanding of how plasticity may affect future adaptation.

## Results

We used a *T. castaneum strain* (Cro1) (Milutinović et al. 2013), which was collected from a wild population in 2010 and adapted to standard control conditions (33 °C, 70% relative humidity (r.h.)) since then. We measured the number of adult offspring as well as gene expression changes relative to control conditions when the beetles were exposed to a drought, a heat and a combined heat-drought treatment (conditions: Dry: 33°C, 30% r.h.; Hot: 37 °C, 70 % r.h; Hot-Dry: 37°C, 30% r.h.). Individuals were transferred to treatments in the egg stage and stayed there during their whole life time. Fitness and expression were measured in females at the age of eleven weeks. Gene expression was measured by RNA-seq using mRNA of the whole body.

### Fitness assay

Offspring number of reproducing females decreased with increasing temperature (F_1,5157_ = 1981.07, P < 2.2e-16) and decreasing humidity (F_1,5184_ = 262.05, P < 2.2e-16), with a stronger effect of heat (−15.98 ± 0.57 SE) than of drought (−5.12 ± 0.50 SE). The lowest offspring number was found when heat and drought were combined (Fig. 1B). Interaction between temperature and humidity was also significant (F_1,5128_= 8.37, P = 0.003835) and led to an additional decrease of −2.22 ± 0.77 compared to purely additive effects. The proportion of reproducing females was significantly different between conditions (χ2= 627.35, df=3, P< 2.2e-16). The highest proportion of non-reproducing females was found in Hot (Supplementary Material S1).

**Figure 1.**
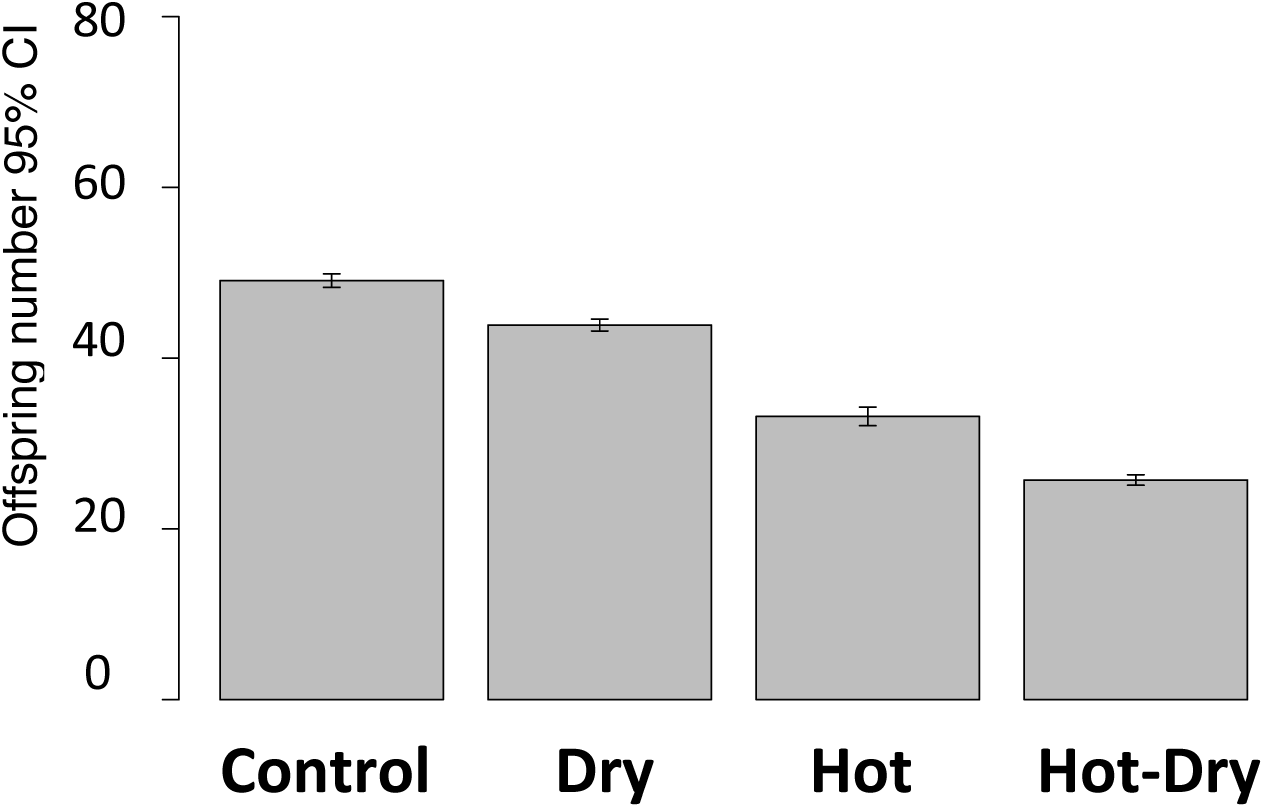
Offspring number of reproducing females in four different conditions: Control (33°C, 70 % r. h.), Dry (33°C, 30 % r. h.), Hot (37°C, 70% r. h.), Hot-Dry (37°C, 30% r. h.) Females could lay eggs for one week and adult offspring was counted five weeks later.

### Gene expression response to heat is stronger than to drought

The number of differentially expressed (DE) genes relative to Control was lowest in Dry and largest in Hot (Fig. 2, Tab. 2). Drought induced up-regulation of 52 and down-regulation of 48 genes. In contrast, the response to heat showed a significantly higher number of DE genes than in Dry (up: 1594, down: 1255; permutation test *P* < 0.004). Overlap between heat and drought responses was significantly higher than expected by chance (χ2= 17.75, d.f.= 1, p-value = 2.516e-05) and included 26 genes (Fig. 2) with responses in the same direction (up: 25, down: 1) and 17 genes with responses in opposite direction. To investigate whether the DE genes were involved in specific biological processes, we performed pathway, protein domains, and Gene Ontology (GO) enrichment tests. Because of a low number of DE genes in Dry, only few enrichments could be detected in Dry (Supplementary Material S2), with up-regulated genes enriched in active ion transmembrane transporter activity (GO:0022853), and down-regulated genes enriched in hydrolase activity, hydrolyzing O-glycosyl compounds (GO:0004553), and protein family Thaumatin (IPR001938). Analysis of the *Tribolium* genome had revealed a high number of genes thought to be involved in endocrine regulation of diuresis, including several that encode putative neuroendocrine peptides like antidiuretic factors (Aikins et al. 2008; Hauser et al. 2008; Li et al. 2008; Park et al. 2008). Nevertheless, none of these genes was found to respond to the Dry treatment.

**Figure 2.**
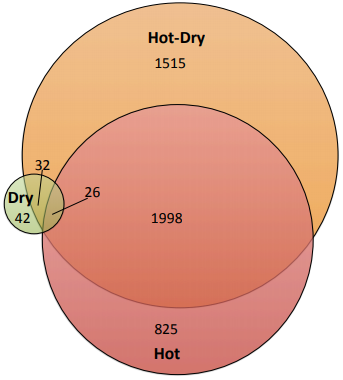
Venn-Diagram showing the number of differentially expressed genes in three treatments relative to control conditions. Overlapping regions represent genes that were found in more than one treatment and changed expression levels in the same direction. Sizes of circles as well as of overlapping regions are proportional to number of genes.

**Table 1:** Results of the fitness assay. Number of adult offspring that females produced within one week of egg-laying in different conditions (Control: 33°C, 70% relative humidity (r.h.).; Dry: 33°C, 30% r.h.; Hot: 37°C, 70% r.h.; Hot-Dry: 37°C, 70% r.h.). For calculating offspring number per female only reproducing females were used

| Condition | N | Females with offspring | Offspring per reproducing female (±SE) |
| --- | --- | --- | --- |
| Control | 1575 | 1514 | 48.99 ± 0.44 |
| Dry | 1642 | 1603 | 43.87 ± 0.43 |
| Hot | 1401 | 1005 | 33.01 ± 0.52 |
| Hot-Dry | 1567 | 1396 | 25.68 ± 0.45 |

**Table 2:** Number of differently expressed genes (FDR < 5%) when comparing different conditions. Positive: higher expression in second condition. Negative: lower expression in second condition. Differential expression analysis was conducted in edgeR (Robinson et al. 2010).

|  | positive | negative | total |
| --- | --- | --- | --- |
| **Dry vs Control** | 52 | 48 | 100 |
| **Hot vs Control** | 1594 | 1255 | 2849 |
| **Hot-Dry vs Control** | 1866 | 1705 | 3571 |
| **Hot vs Dry** | 1553 | 1290 | 2843 |
| **Hot-Dry vs Dry** | 1928 | 1813 | 3741 |
| **Hot-Dry vs Hot** | 101 | 164 | 265 |

Genes up-regulated in Hot were enriched in many metabolic processes, e.g. carbohydrate metabolic process (GO:0005975), Citrate cycle (KEGG 00020), and Pyruvate metabolism (KEGG 00620) (Supplementary Material S1). The most strongly enriched category was chitin metabolic process (GO:0006030). A protein domain analysis also showed significant enrichment of heat shock proteins (IPR031107, IPR018181, IPR008978) (Supplementary Material S2). Down-regulated genes were enriched in pathways for DNA replication (KEGG 03030), nucleotide excision repair (KEGG 03420) and Ubiquitin mediated proteolysis (KEGG 04120) (Supplementary Material S2).

The significant overlap between heat and drought response suggests that these genes are involved in a general stress response. However, no significant functional enrichment could be detected.

### Response to stressor combination is dominated by heat response, single stress responses are mainly not modified in combination

When both stressors were combined in Hot-Dry, we found 3571 DE genes (up: 1866, down: 1705). Among them, 1515 (42.4%) were not found in single stressor treatments. However, only 69 of those genes (up: 30, down: 39) were found significantly DE between Hot-Dry and Dry, or Hot-Dry and Hot. This indicates that in most cases the combined stress did not induce expression changes in a different set of genes, but modified their expression levels over and above their responses to single stressors. Compared to Hot, the Hot-Dry response had a significantly higher magnitude of expression change (permutation test: *P* < 0.0001), a higher number of down-regulated genes (*P* = 0.007), but a similar number of up-regulated and total number of DE genes (*P* = 0.29 and, *P* = 0.054, respectively). The functional response to Hot-Dry resembles the response to Hot, but more enriched GO categories and pathways could be found (see Supplementary Material S2).

To further examine how single stress responses are modified during combination, we classified the DE genes of all treatments into different response categories following Rasmussen et al. (2013) (see Methods and Fig.3). Only 5% of the genes showed a *similar* response mode, with same response to Dry, Hot and Hot-Dry (Fig. 3). Most responding genes (63 %) were classified as *independent*, with a response that is not altered in presence of a second stressor. Most of those genes showed the same response in Hot and Hot-Dry (60.1 % of all genes, Supplementary Material S4), but no response in Dry, in agreement with our DE analysis. 14% had a *combinatorial* response mode: They did not respond to heat and drought alone, but to their combination. These represent cases, in which presence of an additional stressor magnifies the effect of another. Especially interesting are genes with opposite responses to both stressors, but with one response *prioritized* when stressors occur simultaneously. These genes can be indicative of physiological trade-offs that constrain responses to stress combination. We found 8 % of DE genes falling into that category. Most of them showed prioritization of the Hot response in Hot-Dry (7.5 %, Supplementary Material S4). This is in agreement with our DE analysis, which showed a high similarity between responses to Hot and Hot-Dry. 9.5 % of expression responses were classified as *cancelled,* i.e. response disappears when another stressor is added. Most of these genes (6.7 %) showed a significant response in Hot, but not to Dry and returned to control levels in Hot-Dry.

**Figure 3:**
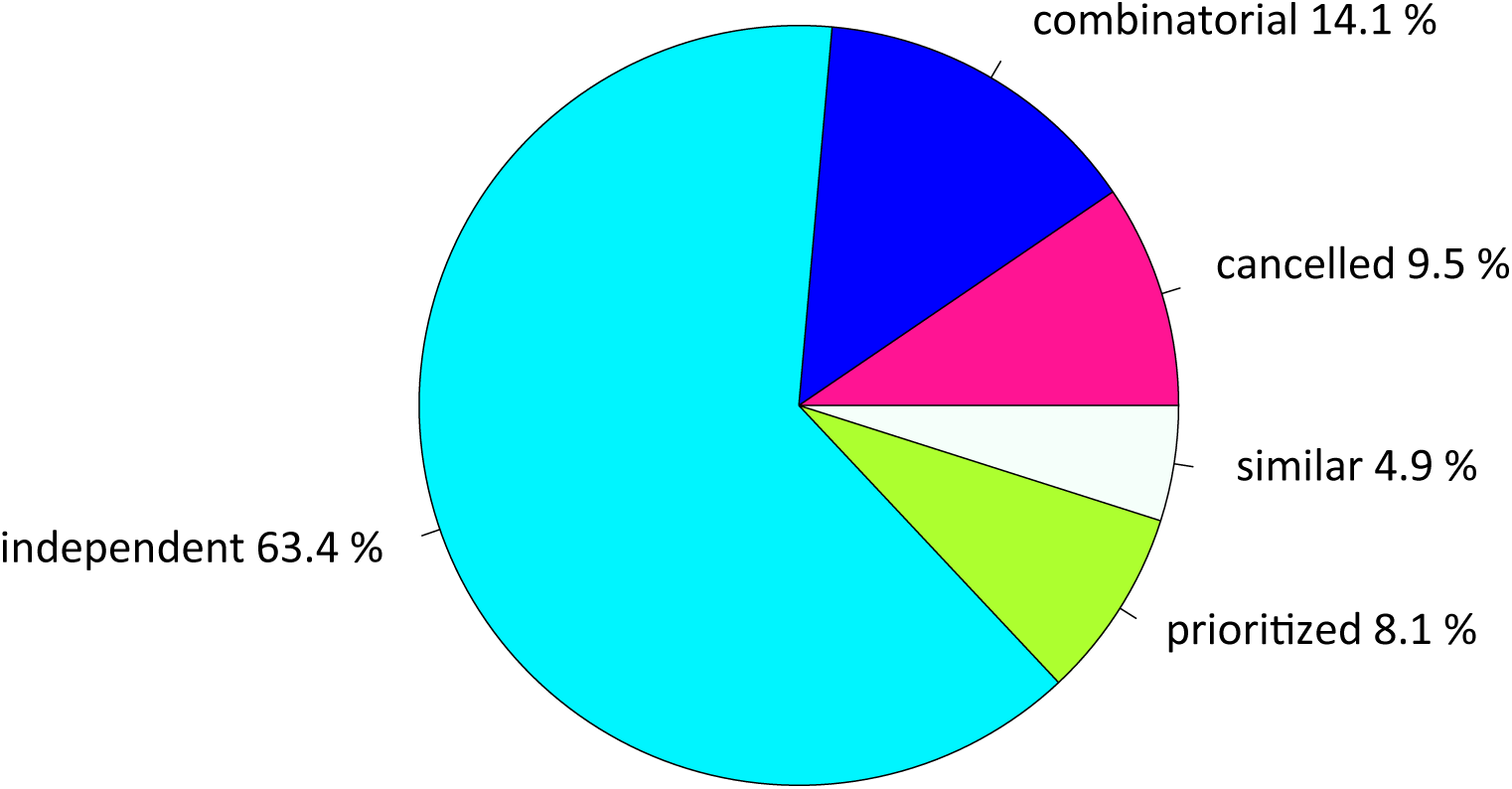
Response modes of genes, which showed significant responses to at least one of the treatments (Dry, Hot, Hot-Dry). *Combinatorial*: Similar levels in the two individual stresses but a different response to combined stresses; *cancelled*: transcript responses to either or both individual stresses returned to control levels; *prioritized*: opposing responses to the individual stresses and one stress response prioritized in response to combined stresses; *independent*: response to only one single stress and a similar response to combined stresses; *similar*: similar responses to both individual stresses and to combined stresses. Subcategories of different response modes, with more details about the most prevalent patterns, are given in supplementary material (S4).

### Immune response: Interaction between drought and heat

We noticed that among the genes up-regulated in Hot, many are involved in immune response (Oppert et al. 2012; Behrens et al. 2014). These transcripts encode putative chitin-interacting proteins (binding, deacetylase), allergens, thaumatins, glucosidases, and serine proteases, whereas these genes are down-regulated at low humidity. For instance, Thaumatin protein family, which is involved in antimicrobial response (Altincicek et al. 2008) showed the strongest enrichment in down-regulated genes (Supplementary Material S2) under drought, and a pathogenesis related protein and several antimicrobial peptides are among those genes that showed the strongest up-regulation in Hot, but were significantly down-regulated in Dry. We applied a gene set enrichment analysis to test for an overrepresentation of immune genes in responses to the different treatments. To avoid including immune genes, which could also be involved in heat stress response (Behrens et al. 2014), we used a test set of 15 antimicrobial peptides, which should be specific to a pathogen response, based on Zou et al. 2007, Oppert et al. 2012, Altincicek et al. 2013 (Supplementary Material S3A). We found that antimicrobial peptides were significantly up-regulated in Hot (p-value = 0.001) and down-regulated in Dry (p-value = 0.001) (Tab. 3). When we compared Hot-Dry with Hot, these genes became down-regulated (p-value = 0.002), indicating that it is mainly the combination of high temperature with relatively high humidity (i.e. 70% r.h.) that activates this response, pointing to a possible higher parasite load under these conditions.

**Table 3.** Results of enrichment analysis of antimicrobial peptides. Gene set enrichment test was conducted in edgeR (Robinson et al. 2010). Prop.Down and Prop.Up give the proportion of genes that are down- and up-regulated. The direction of change is determined from the significance of changes in each direction, and is shown in the Direction column. The p-value provides evidence for whether the majority of genes in the set are DE in the specified direction

| Contrast | Prop. Down | Prop. Up | Direction | p-value |
| --- | --- | --- | --- | --- |
| Dry-Control | 0.5333 | 0 | Down | 0.001 |
| Hot-Control | 0 | 0.4667 | Up | 0.001 |
| Hot-Dry-Control | 0.1333 | 0.2667 | Up | 0.832 |
| Hot-Dry-Hot | 0.4 | 0.0667 | Down | 0.002 |

### Treatment effects on reproduction related processes

To get further insights into the molecular processes that link the observed decline in offspring number with transcriptomic data, we focused on the juvenile hormone (JH), 20-hydroxecdysone (20 E), and insulin/insulin-like peptides (IIS) target of rapamycin (TOR) signaling pathways (IIS-TOR), which are important mediators in the trade-off controlling stress response and reproduction (Schwenke et al. 2016), along with vitellogenin, which is the main nutrient source of eggs, and vitellogenin receptors. We found that heat and heat-drought stress led to a significant down-regulation of all pathways and repression of vitellogenin and vitellogenin receptors (Supplementary Material S3B). Drought did not show any significant effect. A gene set test based on a selection of key reproduction genes within these pathways (Supplementary Material S3C) confirmed that the reproduction genes were mainly down-regulated in Hot and Hot-Dry (Tab. 4).

**Table 4:** Results of gene set enrichment analysis of genes involved in reproduction. Gene set enrichment test was conducted in edgeR using the *roast* function (Robinson et al. 2010). Prop.Down and Prop.Up give the proportion of genes that are down- and up-regulated. The direction of change is determined from the significance of changes in each direction, and is shown in the Direction column. The *P*-value provides evidence for whether the majority of genes in the set are DE in the specified direction. The genes (N=56) were selected based on (Parthasarathy, Sheng, et al. 2010; Parthasarathy, Sun, et al. 2010; Xu et al. 2010; Parthasarathy and Palli 2011).

| Contrast | Prop. Down | Prop. Up | Direction | p-value |
| --- | --- | --- | --- | --- |
| Dry-Control | 0.054 | 0.036 | Up | 0.912 |
| Hot-Control | 0.464 | 0.071 | Down | 0.001 |
| Hot-Dry-Control | 0.589 | 0.071 | Down | 0.001 |

### Selection on expression levels is environment specific

Since we measured offspring number and transcription within the same individuals, we could estimate selection intensity on single gene expression levels in each condition separately by performing a linear regression of relative fitness on standardized expression levels (z-score). In control conditions, expression levels of 2179 genes showed a significant correlation with offspring number (negative: 2158, positive: 21). The two genes under strongest positive selection coded for vitellogenin. Another positively selected gene coded for a serine protease (TC000870) and is involved in oocyte development (GO:0048599). In Dry, Hot and Hot-Dry we could not detect any significant selection on single gene expression levels after correcting p-values for multiple comparisons. To gain a better overview of the variation in selection pressures among treatments and to avoid stringent significance thresholds on single-gene fitness-expression correlations, we then compared selection intensities on all genes between conditions. We found significant negative correlations of selection intensities between Control and all stress treatments, and positive correlations among stress treatments (p-values < 2.2e-16) (Fig 4). Control and Dry had the strongest negative correlation (−0.24), while Hot and Hot-Dry had the highest positive correlation (0.34). Furthermore, significantly DE genes responding to Hot and Hot-Dry were overrepresented among those that were negatively selected in control conditions (Hot: χ2= 158.62, df = 1, p-value < 2.2e-16, Hot-Dry: χ2= 177.97, df = 1, p-value < 2.2e-16), with 361 (22.6 %) up-regulated genes in Hot and 417 (22.3 %) in Hot-Dry. The magnitudes of selection intensities were also significantly different between conditions (median (and SD) of absolute values in Control: 0.09 (0.06), Dry: 0.03 (0.03), Hot: 0.09 (0.07), Hot-Dry: 0.05 (0.04); Kruskal-Wallis rank sum test: χ2= 9333.3, df = 3, p-value < 2.2e-16), with the majority of genes negatively selected in Control, and similar proportions of genes under positive and negative selection in stress treatments (Fig. 5). Estimated selection intensities were rather low compared to those that were reported for morphological and life-history traits in natural populations (Kingsolver et al. 2001; Hereford et al. 2004), but within the range that was estimated for selection on expression levels (McCairns and Bernatchez 2009). Although the Dry treatment had the largest negative correlation with Control expression levels and largest number of genes switching sign relative to Control (9591), it had the lowest magnitude of change in selection intensity of those genes (median: 0.14). In contrast, the Hot environment had the strongest magnitude of change in selection intensity compared to Control (median: 0.19, 6644 genes) followed by Hot-Dry (median: 0.16, 7412 genes). Overall, a greater proportion of genes switched from negative to positive selection (Dry: 0.86, Hot: 0.76, Hot-Dry: 0.8).

**Figure 4:**
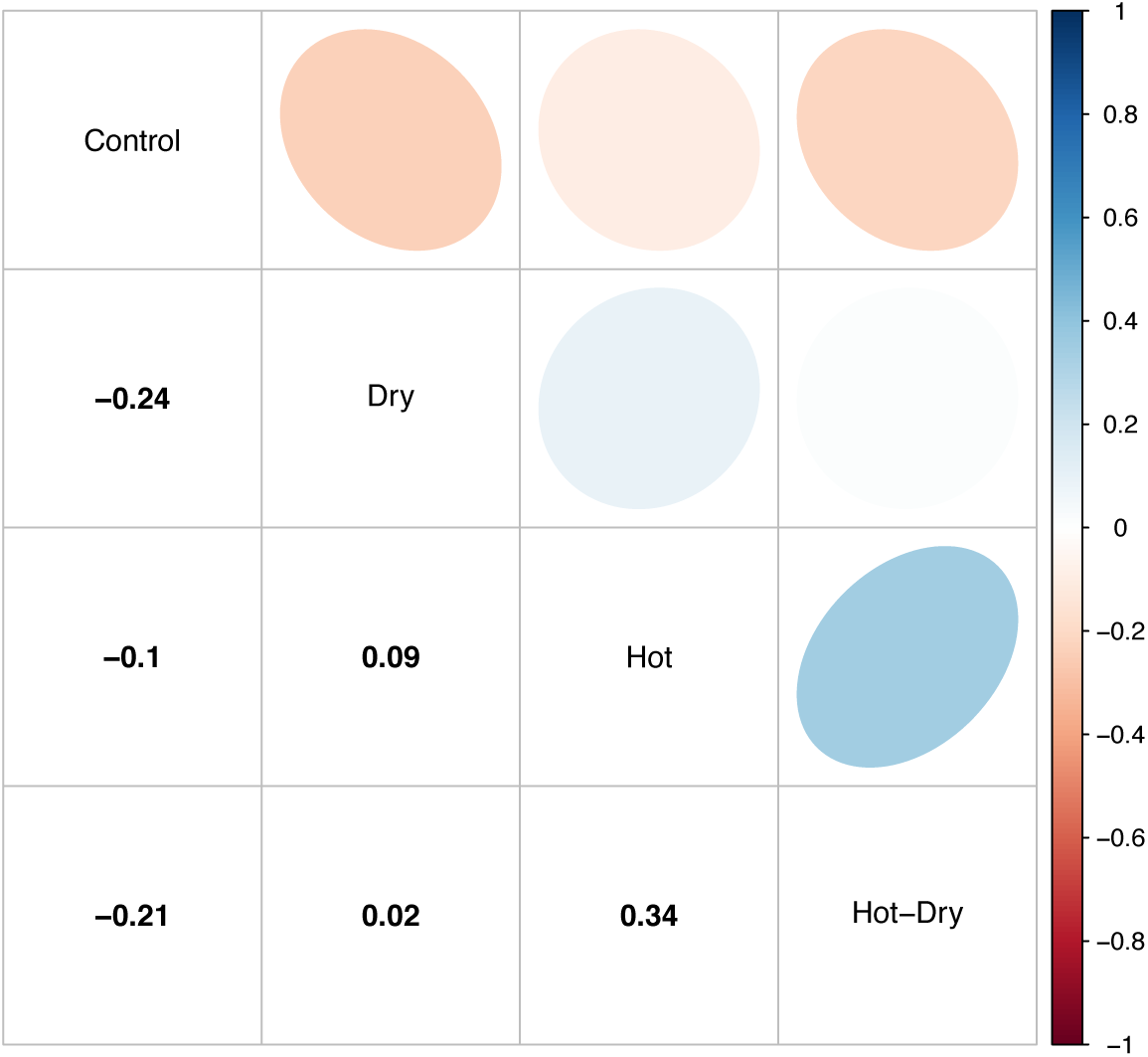
Pairwise correlations of selection intensities on single gene expression levels in different conditions. Blue indicates a positive and red a negative correlation. Values are given in the lower triangle. Confidence intervals for correlations: Control-Dry: -0.25,-0.22; Control-Hot: - 0.11, -0.08; Control–Hot-Dry: -0.23, -0.20; Dry-Hot: 0.08, 0.11; Dry–Hot-Dry: 0.00, 0.03; Hot–Hot-Dry: 0.33, 0.36.

**Figure 5:**
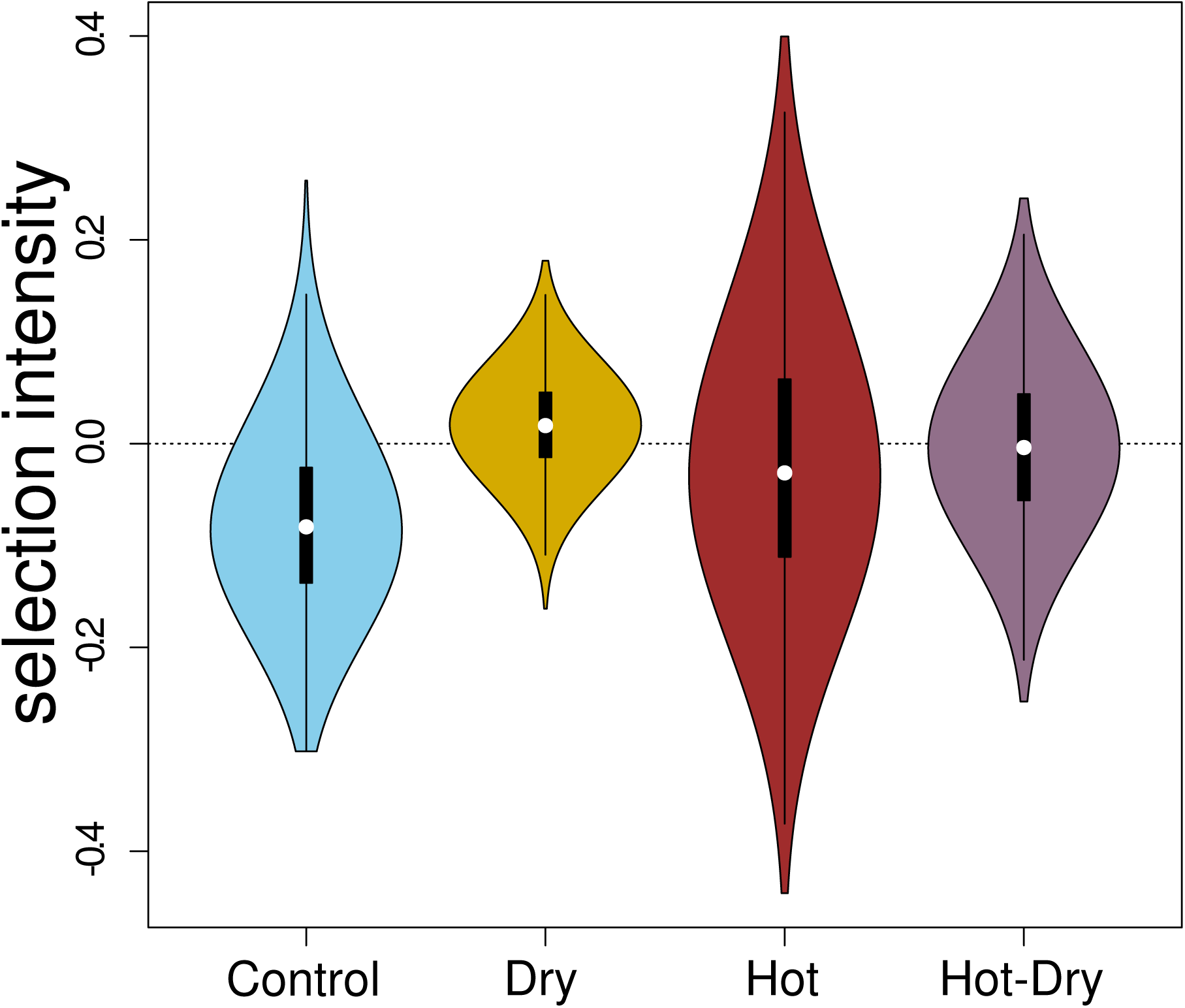
Distribution of selection intensities on gene expression levels under Control and treatment conditions. Each violin plot contains a boxplot of the data. White dots are medians and black rectangles represent inter-quartiles. The selection intensities were obtained as linear regression coefficients of relative fitness (number of adult offspring) on normalized RNA-seq read counts (z-score).

### Is response in gene expression adaptive?

To examine whether the plastic responses in gene expression are adaptive, we tested whether significantly up-regulated genes were more positively selected and significantly down-regulated genes more negatively selected than non-responding genes. We found that the response to Hot-Dry was mainly adaptive: Down-regulated genes were under significantly more negative selection and up-regulated genes under more positive selection compared to non-responding genes, respectively (permutation tests: *P* < 0.0001) (Fig. 6). In contrast, some parts of the response in Dry seemed maladaptive: Down-regulated genes were not under significantly different selection, but up-regulated genes were more negatively selected (Fig. 6). In Hot, the response is partly adaptive since down-regulated genes were significantly more negatively selected, while up-regulated genes were not under significantly different selection (Fig. 6).

**Figure 6:**
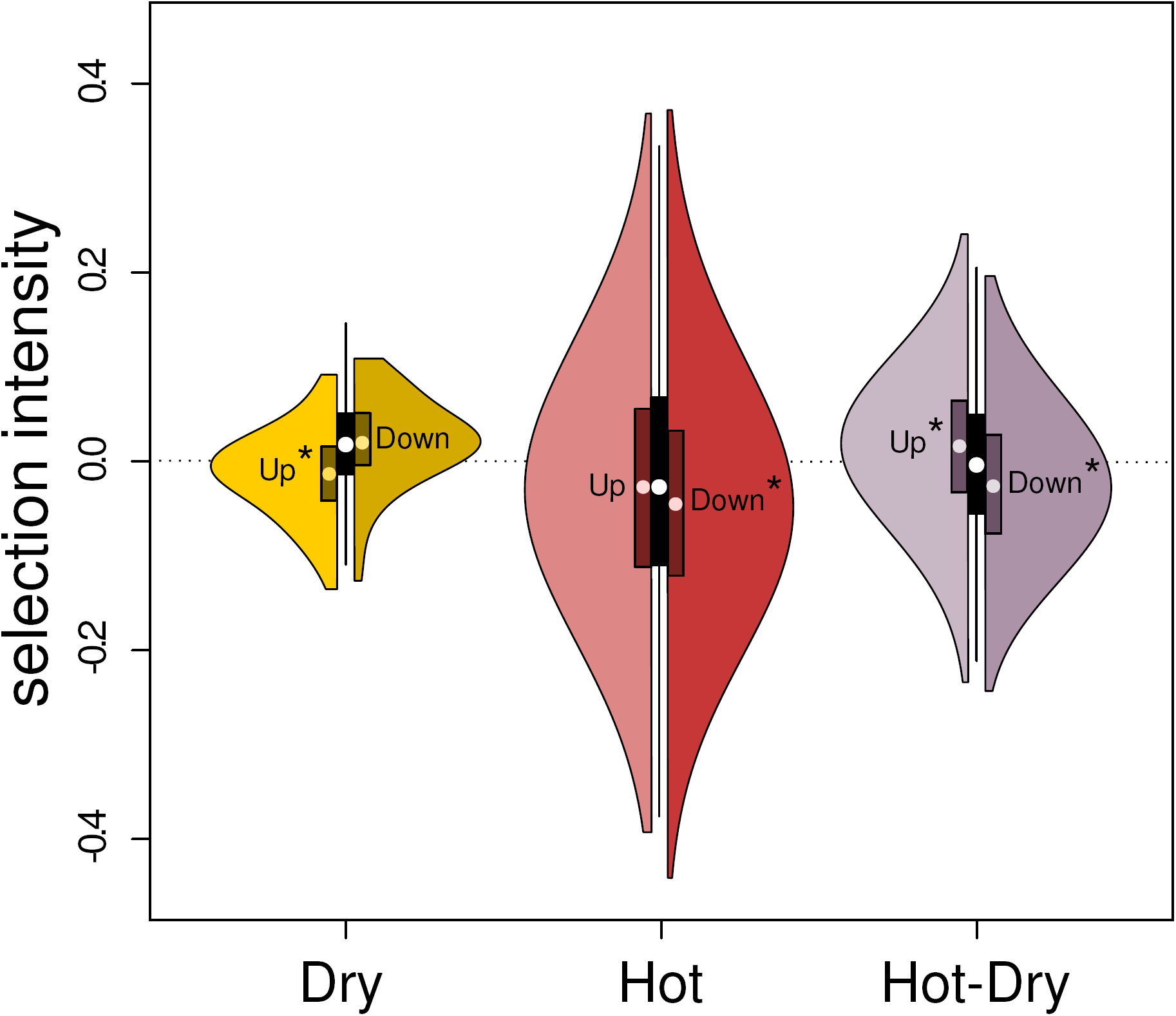
Selection intensities on expression levels of genes that showed significant responses to stress treatments (DE genes). The left half of each violin plot is the distribution of selection intensities of up-regulated genes, relative to Control, while the right half is for down-regulated genes. The median and inter-quartile range are represented as a white dot and a dark rectangle, respectively. The central boxplot represents variation of selection intensities of genes whose expression was not significantly different from expression in Control (at 5% FDR). Significance of the shift in selection intensities of the up- and down-regulated DE genes relative to the non-DE genes is marked with the star symbol (*). Significance was determined with a permutation test with 10,000 permutations (see Methods).

## Discussion

### Selection on gene expression

By measuring gene expression and one component of fitness within the same individuals, we could assess the strength of selection acting on gene expression variation within each environment. The distribution of selection intensities, measuring the change in fitness relative to a change of the phenotypes under selection (Lande and Arnold 1983; Falconer and MacKay 1996), was informative of the type of selection at play in each environment. Under Control conditions, to which the beetles were best adapted, the distribution of selection intensities had the most negative mean and median with a small proportion of transcripts under significant positive selection. This pattern is strongly suggestive of selection against costly gene expression, and thus of some form of purifying selection against non-adaptive variation in expression levels. Gene expression is known to be costly (Castillo-Davis et al. 2002; Wagner 2005; Lang et al. 2009; Frumkin et al. 2017), and selection should therefore favor low expression levels under those mild laboratory conditions. Genes under strongest positive selection were vitellogenin genes playing an important role in reproduction. Vitellogenin genes are also among the most highly expressed genes in the transcriptome. Their strong correlation with offspring number in Control suggests that variation in this component of fitness is mainly driven by differences in egg production among individuals. We could confirm that our measure of fitness is strongly related to fecundity (see Supplementary Material S5), which also confirms that our experimental design was able to capture meaningful correlation of gene expression with reproductive output. Second, shifts in the environment caused an increase of the proportion of beneficial trait variation, with higher median and mean selection intensities in the stress treatments. The changes of the direction of selection, from mostly negative to more positive, affected both the genes that significantly changed their expression plastically (DE genes), and those that did not. The protection provided by stress responses might thus have outweighed the costs of expression seen previously in Control conditions, especially since the direction of selection switched to positive in 80% of the cases. This led to strong negative correlations with selection intensities in Control and suggests that the physiological stress imposed by the stress treatments modified the optimum expression level of a majority of genes in the genome.

In Dry, and to some extent in Hot-Dry, the selection intensities were weaker with a distribution more centered on zero. This may be indicative of stabilizing selection on gene expression levels. Yet, beetles in all three stressful conditions incurred a significant decrease in fitness, and not all plastic changes in gene expression were adaptive. That is, not all were in the direction that increased fitness. The immediate stress responses were thus likely still costly because of trade-offs between stress resistance mechanisms and reproduction, or trade-offs in stress responses between stressors.

### Physiological trade-off between stress response and reproduction within treatments

The heat-induced expression changes were characteristic of stress responses: Up-regulation of heat shock proteins and metabolic pathways, and repression of genes involved in cell growth, cell cycle, and reproduction (Gasch et al. 2000; Kültz 2005; Kassahn et al. 2009; Enders et al. 2015). This pattern of change in gene expression, and the overall reduction of females’ reproductive output, suggests a shift of resource utilization from growth to stress protection. This would be indicative of a physiological trade-off between stress response and reproduction (Salmon et al. 2001; Fedorka et al. 2004; Harshman and Zera 2007). We could confirm that this trade-off was at play because the decrease of females’ reproductive output was mainly driven by a loss of fecundity (egg number) in the Hot and Hot-Dry treatments and not a decrease of larvae survival (see Supplementary Material S5). This matches well with the down-regulation of vitellogenin and other reproduction-related genes (JH, the IIS-Tor and 20E signaling pathways) in Hot and Hot-Dry. Those genes are indeed expected to play an important role in mediating the trade-offs between stress-responses and cellular growth and maintenance (Flatt et al. 2005; Gruntenko and Rauschenbach 2008; Schwenke et al. 2016). Reduction in fecundity was also likely affected by the expression of heat shock proteins, whose fitness costs are known (Feder and Hoffman 1999; Silbermann and Tatar 2000; Kassahn et al. 2009). Finally, a trade-off between immune response and reproduction may also have affected our results. We detected an up-regulation of immune response genes in the Hot treatment, which might be indicative of a higher pathogen load. Alternatively, it might be an indirect response induced by hot and humid conditions, which has evolved in adaptation to pathogens, and might also occur in absence of parasites. Since we did not measure pathogen load, we were not able to distinguish between both. However, a high proportion of non-reproducing females in Hot (Supplementary Material S1) may suggest infections.

### No physiological trade-offs between single stressors

The transcriptomic responses to heat and drought were very contrasted with a small but significant overlap when taken separately. Only a minority of overlapping genes (17 out of 43) showed a trade-off in expression between the two conditions, and no functional enrichment could be detected. Different physiological processes are thus likely affected by the two stresses. This lack of a strong trade-off between the individual stress responses was also evident when investigating the combined transcriptomic response in the Hot-Dry treatment. Most DE genes in Hot-Dry responded to a single stressor independently of the presence of the second stressor (Fig. 3). Such an additive combined stress response can be expected when individual stressors require different protection mechanisms and affect different pathways, and are thus likely not interfering with one another (Folt and Chen 1999). Furthermore, as expected from the single stress responses, the combined response was dominated by the heat response. This is in agreement with many studies showing that heat is a major driver of expression change, especially in ectotherms (Neven 2000; Nguyen et al. 2009; Levine et al. 2011; Chen and Stillman 2012; Morris et al. 2014). We also detected an additive combined effect of heat and drought on fitness, where the average reproductive output in Hot-Dry was close to the combined reduction of fitness in Hot and Dry. Nevertheless, and thanks to our large sample size, we could still detect a significant interaction between heat and drought on offspring number, although the effect was small (-2.22 ± 0.77). Therefore, the addition of a drought stress to the heat stress apparently emphasized the effect of the heat stress, as seen in the transcriptomic response where the Hot-Dry response mainly differed in magnitude compared to the Hot response.

### Adaptive value of plastic responses and indications for evolutionary adaptations

Our results suggest that plastic responses in Hot-Dry and down-regulation in Hot were mainly adaptive to short-term exposures. This adaptive plasticity probably depends on whether *T. castaneum* encountered similar stress in the past and evolved adaptive plastic responses. This should especially be the case for temperature protection mechanisms, essential to ectotherms. However, immediate stress responses might not remain beneficial during long-term adaptation to constant high temperature because they trade off with reproduction. Therefore, costly plastic responses may become maladaptive over time and be reversed despite the immediate benefits provided by their protective functions. This might be particularly the case for heat shock proteins, which were the most strongly responding genes in this experiment. Their immediate protective functions are well known (King and MacRae 2015), as well as the reproductive costs of their overexpression (Silbermann and Tatar 2000). Although adaptive, some of the plastic gene expression changes that we observed may thus be reversed in future evolved populations because other resistance mechanisms may arise during long-term evolution (e.g. enzymes, which are more stable at higher temperatures), making stress protection expendable. On the other hand, part of the genes whose response became associated with lower reproductive output (down-regulation of vitellogenin and other reproduction related processes), would be good candidates for future adaptive up-regulation. Their immediate down-regulation, imposed by trade-offs with stress responses, may then also be reversed. Overall, we expect long-term evolution to decrease differentiation of gene expression relative to the ancestral condition (here Control) in those genes involved in strong fitness trade-offs, but to increase differentiation in genes whose response will remain adaptive.

In the most stressful treatment, Hot-Dry, up-regulated genes under positive selection were enriched in many metabolic processes (e.g. aerobic respiration, citrate metabolic process), while down-regulated genes under negative selection showed enrichment in negative regulation of metabolic processes (GO:0009892). Thus, a part of the adaptive response of gene expression seemed to enhance metabolic activity, potentially improving females’ reproductive output. Those changes constitute candidates for future evolutionary divergence relative to the ancestral population. In Hot, only the down-regulation of genes under negative selection reflected adaptive plasticity. Nevertheless, we can expect to see convergence of the evolutionary patterns of long-term adaptation between the Hot and Hot-Dry selection lines because of their similarity in expression responses and correlation in selection intensities. Results in Dry suggest that maladaptive responses were predominant since up-regulated genes were under more negative selection than non-responding genes. However, responses as well as selection intensities in Dry were generally small, which may reflect the fact that *T. castaneum* has special anatomical adaptations to cope with drought (King and Denholm 2014). Modifying gene expression might therefore play only a minor role in adaptation to dry conditions.

Taken together, our results show a mix of responses sometimes in the direction of natural selection and sometimes in an opposite direction. But, more often than not we found evidence that a significant majority of plastic responses is immediately adaptive, especially in the most stressful conditions. Only in the Dry treatment have we found evidence of maladaptive plasticity. As suggested by Ghalambor et al. (2015), genes with maladaptive plastic responses might be those that will show the strongest changes during evolution. Future evolutionary changes in the three treatments may follow a direction predicted by the selection gradients we measured, although this also depends on the existence of additive genetic variation in expression and genetic correlation with fitness (Robertson 1966; Price 1970; Rausher 1992; Morrissey et al. 2010, 2012; Bonnet et al. 2017). In any case, plastic and non-plastic genes may or may not show evolutionary divergence in expression, depending on underlying trade-offs with reproduction or other traits. While we have estimated the direction and strength of phenotypic selection acting on gene expression, we cannot predict evolutionary changes because we did not attempt to estimate the additive genetic co-variance of the traits under selection (estimated with their G-matrix: Lande 1979; Walsh and Blows 2009). Genetic covariation among phenotypic traits and with fitness are tedious to estimate and necessitate much larger sample sizes than we had here. (but see McGraw et al. 2011, Blows et al. 2015 for recent attempts using transcriptomic data). Genetic covariance in expression can be caused by shared regulatory elements (e.g., same pleiotropic transcription factors) and lead to correlated responses to selection among genes, which can strongly deviate from predictions of single-gene selection responses (Lande 1979; Walsh and Blows 2009). Nevertheless, our measures of phenotypic selection allowed us to test if up-regulated or down-regulated genes were targets of positive or negative selection pressures, respectively. We could thus test if plastic expression changes were responding to environmental changes in a direction favored by selection. Future work based on a deeper sampling and long-term evolution will address whether those selection pressures translated into corresponding evolutionary changes as predicted by genetic covariation of expression with fitness.

One drawback of our study is the exclusion of males from the gene expression measurements. We decided to focus on females, because they have to invest more into reproduction compared to males. Energy allocation trade-offs between responses to different stressors, as well as between stress protection and reproduction should be more pronounced here. We could also show that number of adult offspring was mainly dependent on egg number (Supplementary Material S5), suggesting a minor contribution of males to fitness. Correlation of male expression levels with fitness, and hence selection intensity, would probably be lower, because also dependent on the status of mating females.

## Conclusions

Our approach shows how transcriptomics can be used to get information about the relative importance of different stressors, their interaction and the potential constraints acting on plastic and evolutionary responses when several environmental variables change at the same time. We were thus able to evaluate the immediate adaptive value of the plastic changes in gene expression. By adaptive, we here meant immediate fitness advantage or disadvantage associated with changes in gene expression. Therefore, our study strongly contributes to understanding how plasticity affects fitness at the early onset of adaptive divergence and gives indications of potential future changes in gene expression. It shows that some parts of the plastic response are adaptive, whereas others are maladaptive. However, further work is needed to clarify how we can use plastic responses to predict long-term evolutionary outcomes, for instance by using long-term evolution experiments.

## Material and Methods

### Animal rearing and stress treatments

We used the *Tribolium castaneum* Cro1 strain (Milutinović et al. 2013), collected from a wild population in 2010 and adapted to lab standard conditions (33°C, 70% relative humidity) for more than 20 generations. Beetles were kept in 24h dark on organic wheat flour mixed with 10% organic baker’s yeast. We sterilized the flour and yeast by heating them for 12h at 80°C before use. We tested the response of fitness and gene expression of the beetles to heat, drought, and a combination of both stressors. The conditions in the treatments were: Hot: 37°C and 70% r. h., Dry: 33°C and 30% r. h., Hot-Dry: 37°C and 30% r. h. Parents of the experimental beetles were reared and mated in control conditions at the age of four weeks in 15 mL tubes with 1 g of medium. Each virgin male was mated with a virgin female. After four days, in which the beetles could mate and lay eggs, each mating pair was transferred to a new vial. We repeated this three times, resulting in four vials per mating pair containing medium and eggs. Vials of each mating pair were randomly assigned to the four different conditions, resulting in full-sib families split across all conditions. Male and female offspring (four females and four males per family and condition) were separated at the pupal stage and transferred to 10 mL tubes with 1 g of medium and remained there until they were used for the fitness assay eight weeks later. After the fitness assay, males and females were transferred to 1 mL tubes, frozen in liquid nitrogen and stored at -80°C. The fitness assay was started in the morning and stopped in the afternoon one week later by removing the mating pair, which was then immediately frozen. Beetles should not show a diurnal cycle since they were kept in 24h dark.

### Fitness assay

To test the effects of the different conditions on fitness, we measured reproduction in 6183 virgin females (ca. 1500 per condition, Table 1). We mated each virgin female with one unrelated male from the same condition in 15 mL tube with 1 g medium. The male was removed after 24 h. Females were removed from the tubes after one week of egg laying, and 9 g medium was added to provide food for the developing offspring. After five weeks the number of offspring was counted. At this time, all offspring had reached the adult stage. Some females did not produce any offspring, in proportions that differed between conditions. To test whether there was an effect of treatment on the number of reproducing and non-reproducing females, we used a generalized linear mixed model with reproduction success (binomial: offspring/no offspring) as response and condition as fixed effect. Since some of the tested females and males were full-sibs and developed within the same tube, we used male and female families as random factors to account for non-independence due to relatedness and a shared environment during development. To test how offspring number of reproducing females was influenced by conditions we used a linear mixed model with offspring number as response, temperature, humidity and their interaction as fixed effects and female and male family as random factors. Denominator degrees of freedom were estimated using Satterthwaite approximation. Statistical analyses were performed using the Lme4 package (Bates et al. 2015) version 1.1-17 in R (R Core Team 2015).

### RNA extraction, library preparation and sequencing

183 female beetles with known fitness (ca. 45 per condition), which had been stored at -80°C, were homogenized in Tri-Reagent^®^ (Zymo Research, California, USA) using an electric bead mill. RNA was extracted with the RNA Mini Prep kit (Zimo Research, California, USA) following the instructions of the manufacturer. RNA-quality was checked with the Bioanalyzer 2100 (Agilent, Waldbronn, Germany) and concentrations were measured with aQubit^®^ Fluorometer (Life Technologies, California, USA). Libraries were created with 500 ng RNA for each individual separately with the LEXOGEN mRNA-Seq Library Kit following the manual (LEXOGEN GmbH, Vienne, Austria). Library quality was checked on a TapeStation (Agilent, Waldbronn, Germany) and concentrations were determined by qPCR. Libraries were diluted to the same molarity. Concentrations of dilutions were checked again by qPCR and libraries were pooled (36 libraries per pool). All treatments were randomized during RNA-extraction, library preparation, and sequencing. The single-end sequencing was performed in five runs on the Illumina NextSeq 500 (Illumina, Inc, California, USA) using the 75 cycles High Output Kit. Each run resulted in 550-600 million reads that passed the internal sequencer filter. After quality control using FastQC (www.bioinformatics.bbsrc.ac.uk/projects/fastqc), reads were mapped against the reference genome (ftp://ftp.ensemblgenomes.org/pub/release30/metazoa/gtf/tribolium_castaneum/Tribolium_castaneum.Tcas3.30.gtf.gz) with STAR v.2.5 (Dobin et al. 2013) (adaptors were trimmed and the first 10 bases were hard trimmed, minimum average quality Q10, minimum tail quality 10, minimum read length 20). We then used FeatureCounts (Liao et al. 2014) to count the number of reads that mapped to each gene in the reference genome. Mapping as well as read counting was performed within the data analysis framework SUSHI (Hatakeyama et al. 2016).

### Differential expression and enrichment analysis

We conducted a differential expression analysis using the R package edgeR (Robinson et al. 2010). We tested for differently expressed genes between the treatments (Dry, Hot, Hot-Dry) relative to the control as well as to each other. A gene is classified as DE with a FDR ≤5% after adjusting for multiple testing (Benjamini and Hochberg 1995). To test whether the number of DE genes (relative to Control) was significantly different between two environmental conditions a permutation tests was used. For each permutation entire RNA-seq samples of the two groups were randomly assigned to conditions and the edgeR analysis repeated. Significance was assessed by number of times the observed DE number was higher than the DE number obtained by permutations. To test whether the magnitude of change in expression levels relative to control was significantly different between Hot and Hot-Dry, we performed a permutation test. Absolute log2-fold changes of each transcript were randomly assigned to the two groups and differences in the mean were calculated. We then compared the distribution of differences obtained by permutations to the observed difference between mean absolute log2-fold changes in expression. Gene set enrichment analyses for immune response genes and reproduction related genes were conducted in edgeR using the *roast* function (Wu et al. 2010). The significance cutoff for genes contributing to the proportion of down-regulated genes is 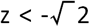 and 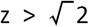 for proportion of up-regulated genes (Robinson et al. 2010). A GO enrichment analysis of DE genes was performed with gProfiler Version: r1622_e84_eg31 (Reimand et al. 2011) and pathway and protein domain enrichment analysis with STRING v.10.0 (Szklarczyk et al. 2015).

### Classification of response mode

Following Rasmussen et al. (2013), we created 20 predefined expression profiles each representing a potential response when two single stressors are combined: *Combinatorial*: similar expression levels in single stress treatments but a different level in stress combination, *cancelled*: response to one or both single stressors but expression levels similar to control conditions when both are combined, *prioritized*: opposite responses to single stressors and expression levels in combination similar to one of them, *independent*: response to only one single stressor and the same response in combination, *similar*: same response in each of the two single stressor treatments, and combination. For creating predefined expression profiles we used 0 as control level, 1 and -1 as expression levels for up- and down-regulation, e.g. expression profile for an independent response could be: CT:0, D:0, H:1, HD:1. We then created a dataset consisting of all genes showing a significant response in at least one treatment (4419 genes in total). Correlation between normalized read counts (cpm, TMM normalization) of these genes and each of the predefined expression profiles was tested and genes were assigned to the category with the highest correlation.

### Selection

We measured selection intensity on gene expression separately for each treatment using univariate linear regression methods (Lande and Arnold 1983; Brodie III et al. 1995). Fitness (number of adult offspring) was normalized by dividing each individual value by the mean (w’ = w_i_/mean(w)). For each gene, expression levels were first normalized to cpm (counts per million, TMM normalization) using edgeR and then transformed to standardized z-scores by subtracting the mean and dividing by the standard deviation (z = (x_i_-mean(x))/ SD(x)). Resulting regression coefficients of relative fitness on standardized expression levels give an estimate of the selection intensity. P-values were corrected for multiple comparisons. To test whether up- and down-regulated genes were under significantly different selection compared to genes without a significant response, we used permutation tests. For each permutation (10,000 for each test) we randomly assigned the categories “not DE” and “up” (or “down” respectively) to each estimated selection intensity and calculated the difference in mean selection intensity between both groups. Significance was tested by counting the number of permutations that showed a difference higher or equal to the observed one.

## Supporting information

## Acknowledgements

This work was supported by the Swiss National Science Foundation, grants PP00P3_1144846 and PP00P3_176965 to FG. We thank Sonja Sbilordo, Julian Bauer, Valérian Zeender for helping during the fitness assay, Maria Domenica Moccia for her help during the fitness assay and RNA extraction and Lucy Poveda for advice in library preparation and sequencing. We thank Fabian Freuler, Marcel Preisig and Cátia Pereira, who conducted the fecundity assay presented in the supplementary material S5.

## Notes

#### Summary of Updates

Introduction and Discussion updated to clarify use of phenotypic correlation with fitness to estimate selection gradients on gene expression; Figures 1, 3, 5, and 6 updated. Supplementary files updated

