## Supplementary material for "Combining transcriptomic and fitness data reveals additive and mostly adaptive plastic responses of gene expression to multiple stress in *Tribolium castaneum*"

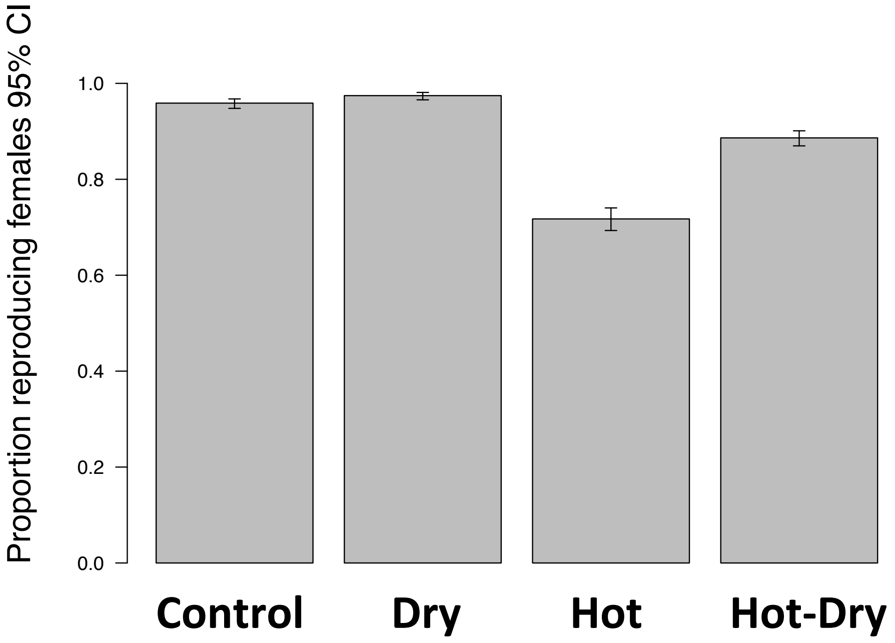


**Figure S1**. Proportion of reproducing females in four different conditions: Control (33°C, 70 % relative humidity), Dry (33°C, 30% r.h.), Hot (37°C, 70% r.h.), Hot-Dry (37°C, 30% r.h.).
