## Supplementary material for "Combining transcriptomic and fitness data reveals additive and mostly adaptive plastic responses of gene expression to multiple stress in *Tribolium castaneum*"

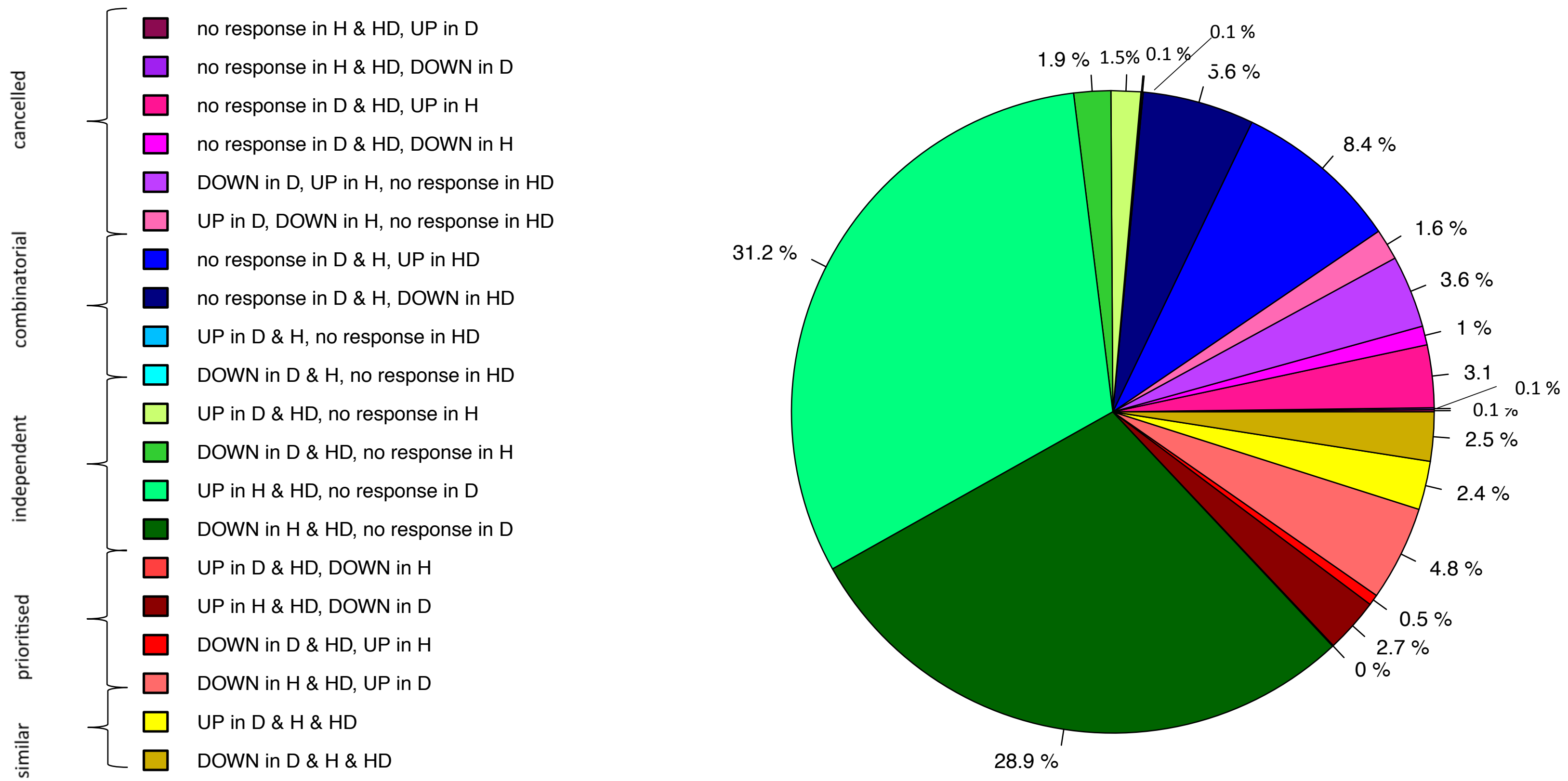


**S4:** Subcategories of different response modes giving more details about the most prevalent patterns : Response modes of significantly responding transcripts in the stress treatments (Dry (D), Hot (H), Hot-Dry (HD). Combinatorial: Similar levels in the two individual stresses but a different response to combined stresses; cancelled: transcript response to either or both individual stresses returned to control levels; prioritized: opposing responses to the individual stresses and one stress response prioritized in response to combined stresses; independent, response to only one single stress and a similar response to combined stresses; similar: similar responses to both individual stresses and to combined stresses.
