## Supplementary material for "Combining transcriptomic and fitness data reveals additive and mostly adaptive plastic responses of gene expression to multiple stress in *Tribolium castaneum*"

During two experiments in 2015 and 2017 we tested for differences in egg number, hatching rate and larvae survival in different conditions. In 2015 one single line was used, in 2017 individuals of seven different replicate lines were tested. The lines were adapted to control conditions and originated from the same strain (Cro1) (Milutinović et al. 2013) that we used for our gene expression/fitness study.

A male and a female beetle could mate for 48 h in a petri dish with 0.5 g of medium. The mating pair was transferred to a new petri dish and stayed there again for 48 h. Eggs were counted after the removal of the mating pair and 2 g of medium added. Larvae were counted after one week and adults five weeks later. We used the sum of eggs, larvae and adults of the two petri dishes for each mating pair for statistical analysis.

To test for significance we used a linear mixed model (Rpackages lme4 (Bates et al. 2015) ) and included line as a random effect. Statistical analyses were performed in R version 1.1-17 (R Core Team 2015).We found that egg number was significantly influenced by condition ($F_{2,214} = 47.87$ , $P < 2.2e-16$). Hot as well as Hot-Dry reduced the number of eggs significantly (Hot: - 19.05 ± 2.43 SE; Hot-Dry: - 21.73 ± 2.51).

There were no differences in larvae survival ($F_{2,210} = 1.43$ , $P = 0.24$), but hatching rate was lower in Hot and in Hot-Dry ($F_{2,212} = 24.32$ , $P = 3.097e-10$). Fig S5 and Tab S5 show that it is mainly the difference in number of laid eggs that is responsible for the differences in number of adult offspring.

**Table S5:** Reproduction in different conditions: Number of eggs that were layed by single females within four days and the resulting larvae and adults. Control: 33°C, 70% r.h.; Hot: 37°C, 70 % r.h.; Hot-Dry: 37°C, 30 % r.h.

| | N | Eggs ± SE | Larvae ± SE | Adults ± SE | Hatching rate ± SE | Larvae survival ± SE |
|---|---|---|---|---|---|---|
| **Control** | 114 | 56.19 ± 4.91 | 48.46 ± 4.49 | 43.34 ± 3.52 | 0.8477 ± 0.0370 | 0.9160 ± 0.0262 |
| **Hot** | 52 | 37.13 ± 5.19 | 27.77 ± 4.84 | 24.22 ± 3.91 | 0.7301 ± 0.0449 | 0.9250 ± 0.0299 |
| **Hot-Dry** | 53 | 34.46 ± 5.08 | 25.05 ± 4.71 | 21.15 ± 3.78 | 0.6001 ± 0.0428 | 0.8894 ± 0.0288 |

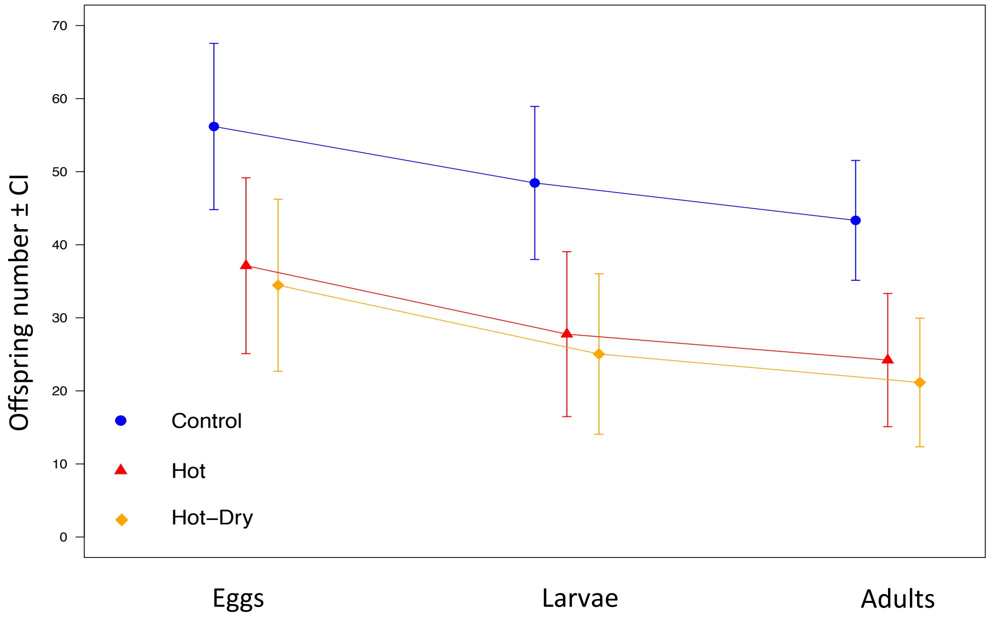


**Figure S5:** Offspring numbers in different developmental stages in different conditions. Lower number of adult offspring in stressful conditions Hot and Hot-Dry is mainly caused by a reduction in eggs. Control: 33°C, 70% r.h, N=114.; Hot: 37°C, 70 % r.h., N=52; Hot-Dry: 37°C, 30 % r.h., N=53.

R Core Team. 2015. *R: A Language and Environment for Statistical Computing*
